# Endoplasmic reticulum-anchored nonstructural proteins drive human astrovirus replication organelle formation

**DOI:** 10.1101/2025.06.25.661519

**Authors:** Brooke Bengert, Samaneh Mehri, Madeline Holliday, Nicholas J. Lennemann

## Abstract

Human astroviruses (HAstV) are major cause of acute, non-bacterial gastroenteritis and have been implicated in severe infections of the nervous system. Despite global prevalence, there are no established treatments for HAstVs due to a lack of understanding of the fundamental biology of infection, including mechanisms of viral replication. Like all positive-stranded RNA (+ssRNA) viruses, infection induces remodeling of host membranes into replication organelles (ROs). However, the intracellular membrane source and viral proteins involved in the coordination of HAstV ROs remain poorly defined. Using immunofluorescence microscopy, we determined that HAstV1 infection drives extensive restructuring of the endoplasmic reticulum (ER) to concentrate RNA replication and virus packaging. Long-term, time-lapse imaging of the ER and time point transmission electron microscopy revealed that temporal manipulation of ER membrane corresponds with the emergence of ER-contiguous double membrane vesicles (DMV). The co-expression of transmembrane nonstructural proteins nsp1a/1 and nsp1a/2 established similar DMV networks in the absence of an active infection. Further, super resolution microscopy revealed the organization of these two viral proteins in RO-like arrangements within the perinuclear region of infected cells. Together, these findings enhance our understanding of HAstV1-induced RO biogenesis, highlighting nsp1a/1 and nsp1a/2 as exploitable targets for the design of antivirals restricting astrovirus replication.

**Significance:** Human astroviruses (HAstV) are understudied, globally prevalent pathogens capable of causing potentially fatal infections of the central nervous system in children, the elderly, and immunocompromised individuals. Despite the capacity to cause devastating disease, there are no established vaccines, antivirals, or therapeutics available combat HAstV infections, in part due to a lack of knowledge for the functions of several viral proteins. This study furthers our understanding of HAstV manipulation of the host cell by identification of the viral proteins responsible for the biogenesis of virus-induced double membrane vesicles, which are a hallmark of replication.

## Introduction

The *Astroviridae* family is composed of small, non-enveloped, positive-sense RNA viruses (+ssRNA) that infect a variety of animal hosts (1, 2). Astroviruses are classified into two genera by host range, including viruses isolated from both mammals (*Mamastrovirus*) and birds (*Avastrovirus*) that cause a variety of mild to severe pathologies (2). Human astroviruses (HAstV) belong to the *Mamastrovirus* genus and are comprised of three genetically distinct groups: the classical serotypes (HAstV1-8) and two divergent lineages, MLB and VA (1, 2).

Classical astroviruses are a leading cause of non-bacterial gastroenteritis, representing 2 to 9% of all acute cases in children across the world and an estimated 3.9 million cases in the U.S alone every year (2, 3). Divergent human astroviruses are associated with extra-gastrointestinal and neurotropic infections in immunocompromised children, which can be fatal (4–9). Despite the potential to cause severe disease, *Astroviridae* remains an understudied viral family of global health importance.

The ∼6.8–7.4 kilobase (kb) astrovirus genome is comprised of three overlapping open reading frames (ORFs) that are translated as polyproteins: ORF1a, ORF1b, and ORF2(1). Furthermore, it has been shown that classical HAstVs and MLBs contain an alternative translation start codon within ORF2 that produces protein X, which acts as viroporin that is important for virion egress (10). Viral nonstructural proteins (nsp) are generated from ORF1a and ORF1b, whereas structural proteins are translated from ORF2 transcripts driven by a subgenomic promoter (SGp) present near the 3’ end of ORF1b (1, 11–13). The nsp1a polyprotein (ORF1a) encodes five proposed nonstructural proteins: a single-pass transmembrane protein (nsp1a/1), a multi-pass transmembrane protein (nsp1a/2), a serine protease (nsp1a/3), a cap-like viral protein linked to the genome (nsp1a/4-VPg), and a phosphoprotein (nsp1a/4-p20) (14–17). Production of the RNA-dependent RNA polymerase (RdRp, nsp1b) occurs through an infrequent (−1) ribosomal frameshift (RFS) at a conserved RNA stem loop at the junction of ORF1a and ORF1b, resulting in co-translation of nsp1ab (18–20). While the genomic organization and predicted protein products of ORF1a and ORF1b have been defined, the specific roles of several astrovirus nonstructural proteins have yet to be characterized, which is critical for understanding the mechanisms of replication and pathogenesis (21).

The viral polyprotein is targeted to the endoplasmic reticulum (ER) by nsp1a/1, which contains a transmembrane domain that acts as a signal peptide for nsp1a/2 (22, 23). Host signal peptidase has been proposed to mediate the cleavage between nsp1a/1 and nsp1a/2 in the ER lumen, while the remaining cytoplasmic junctions between individual nonstructural proteins are cleaved by the viral protease, nsp1a/3 (21, 22, 24, 25). Previous studies have reported that nsp1a/1 and nsp1a/4 colocalize with the ER and replicating RNA (23, 26). Additionally, differing sequences present within the hypervariable region (HVR) of nsp1a/4 have been linked to increased virus recovered from fecal samples, suggesting nsp1a/4 may modulate replication or pathogenicity (27). Still, the molecular functions and interactions of nsp1a/1, nsp1a/2, and nsp1a/4 during infection remain elusive.

Like all +ssRNA viruses, astrovirus infection induces extensive rearrangements of intracellular membranes for productive replication, including the formation of double membrane vesicles (DMV) adjacent to viral particles within infected cells (28–31). DMVs, typically derived from the ER, belong to a broad class of +ssRNA virus-induced replication organelles (ROs) (28). These essential structures are thought to sequester the viral RNA and replication machinery within the same topological compartment, as well as shield double-stranded RNA (dsRNA), an immunostimulatory replication intermediate of +ssRNA viruses, from pattern recognition receptors (PRRs) in the cytoplasm (28, 31, 32). Among other +ssRNA viruses, membrane reorganization is mediated by viral nonstructural proteins with transmembrane or membrane-interacting domains (33–40). However, there is a lack of experimental evidence for the intracellular membrane source and viral proteins that drive astrovirus RO biogenesis.

In this study, we employed a variety of complimentary microscopy techniques to characterize the extensive restructuring of ER membrane that occurs during HAstV1 infection. We report that replication occurs within ER-derived DMVs, and assembly of progeny virus occurs within a unique ER-enclosed membrane boundary. Further, the expression of transmembrane viral proteins nsp1a/1 and nsp1a/2 is sufficient to induce the formation of DMV networks in the absence of an active infection. These findings establish that nsp1a/1 and nsp1a/2 serve as key effectors of ER remodeling and RO biogenesis during astrovirus infection, making them ideal targets of antivirals inhibiting viral replication.

## Results

### HAstV1 infection drives remodeling of ER membrane

Given colocalization of nsp1a/1 and nsp1a/4 with ER membrane, it has been suggested that astrovirus ROs originate from the ER (23, 26, 31). Thus, we sought to confirm remodeling of the ER during HAstV1 infection using immunofluorescence microscopy. Infection of Caco2 cells resulted in aggregation of the ER in regions of dsRNA staining, which mark sites of +ssRNA virus replication (Fig. 1A). Quantification of the total area and size of distinct clusters of ER marker (calnexin) signal between mock and HAstV1 infected cells revealed that infection results in significant ER membrane condensation and network fragmentation, respectively (Fig. 1B).

**Figure 1.**
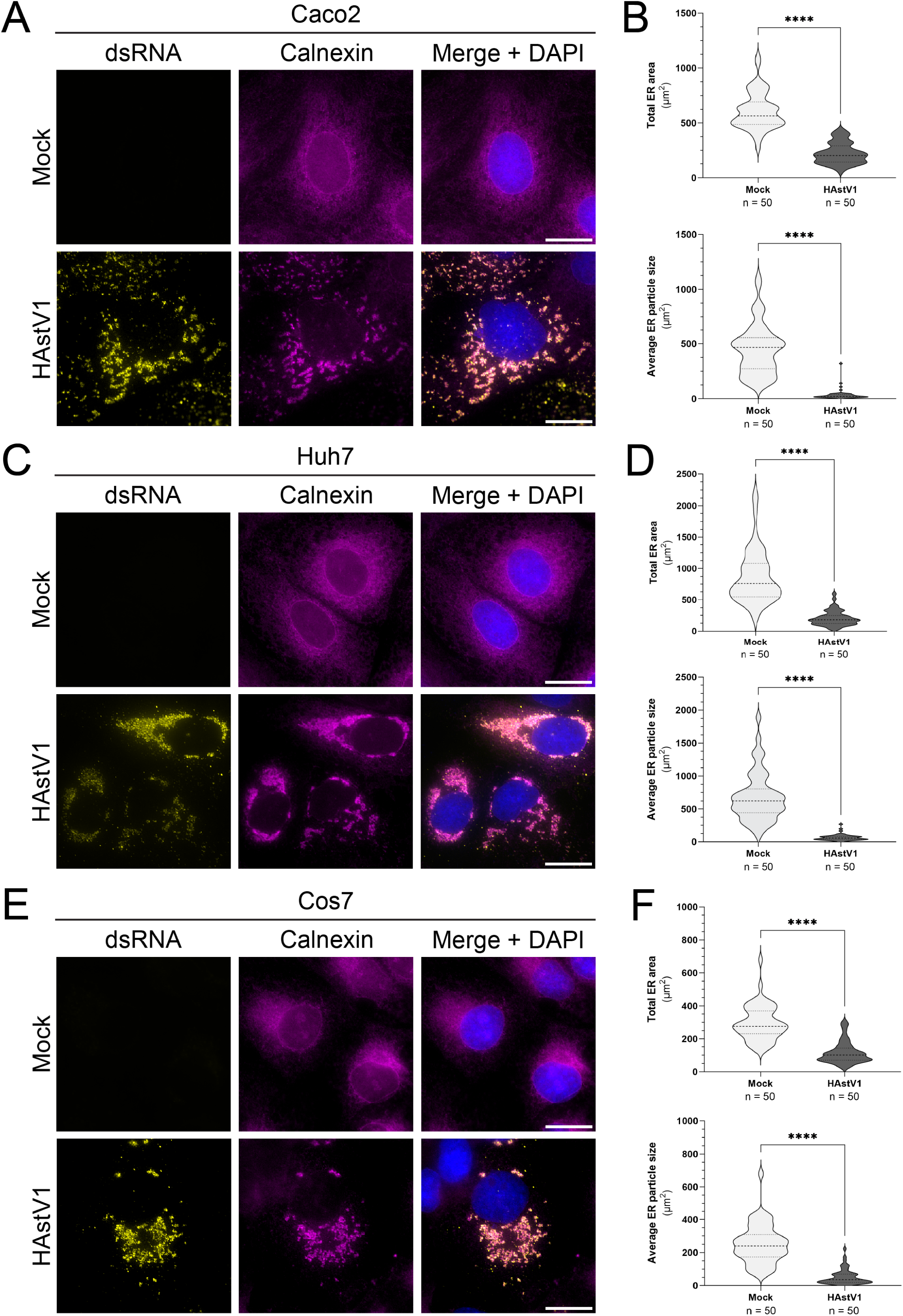
Cell type independent astrovirus ER remodeling. (A, C, E) Representative immunofluorescence staining of mock or HAstV1 infected (MOI = 3; 24 hpi) cells of the indicated type. Scale bars represent 20 µm. (B, D, F) Quantification of the total area of ER signal (top) and mean area of ER particles (bottom) between groups by particle analysis using FIJI ImageJ software and an unpaired t-test (**** p < 0.0001). Thresholding was applied uniformly between groups for n = 50 cells across 3 independent experiments.

Furthermore, HAstV1-driven restructuring of the ER occurs in multiple cell types, as infection of two other cell lines that support HAstV1 replication (Huh7 and Cos7) produced similar aggregates of ER at 24 hours post infection (hpi) (Fig. 1C and 1E). Quantification of the ER morphology in these additional cell types demonstrated the same trend of decreased ER area and particle size (Fig. 1D and 1F). Thus, the consistent remodeling of the ER across multiple permissive cell lines supports a conserved role for ER membrane during astrovirus replication.

### Temporal restructuring of ER membrane correlates with the biogenesis of DMVs

The ER is a dynamic organelle that undergoes continuous remodeling as part of normal cellular function and growth (41, 42). Therefore, we sought to capture temporal changes to the ER by long term, time-lapse imaging of a fluorescent protein localized to the ER lumen, mCherry-KDEL (mCh-KDEL). Notably, fixed IF imaging of HAstV1-infected Huh7 cells resulted in similar clustering of exogenously expressed mCh-KDEL compared to endogenous calnexin staining (Fig. 2A and 1B). Furthermore, live-cell imaging demonstrated that astrovirus infection-dependent ER manipulation progresses through distinct membrane intermediates throughout the course of a 24-hour infection. Fragmentation of the ER network begins at 12–14 hpi, intensifies by 18 hpi, and culminates with the condensing of ER aggregates at the perinuclear space by 20 and 24 hpi (Fig. 2B, Supplemental Movie 1 and 2). Closer examination of the images acquired every 20 minutes between 14–18 hpi confirmed that ER aggregates likely form due to recruitment and compacting of peripheral ER rather than membrane proliferation (Fig. 2C, Supplemental Movie 3). Indeed, we found no difference in the abundance of several ER proteins (calnexin, CLIMP63, RTN4) between mock and infected cell lysates (Fig. S1).

**Figure 2.**
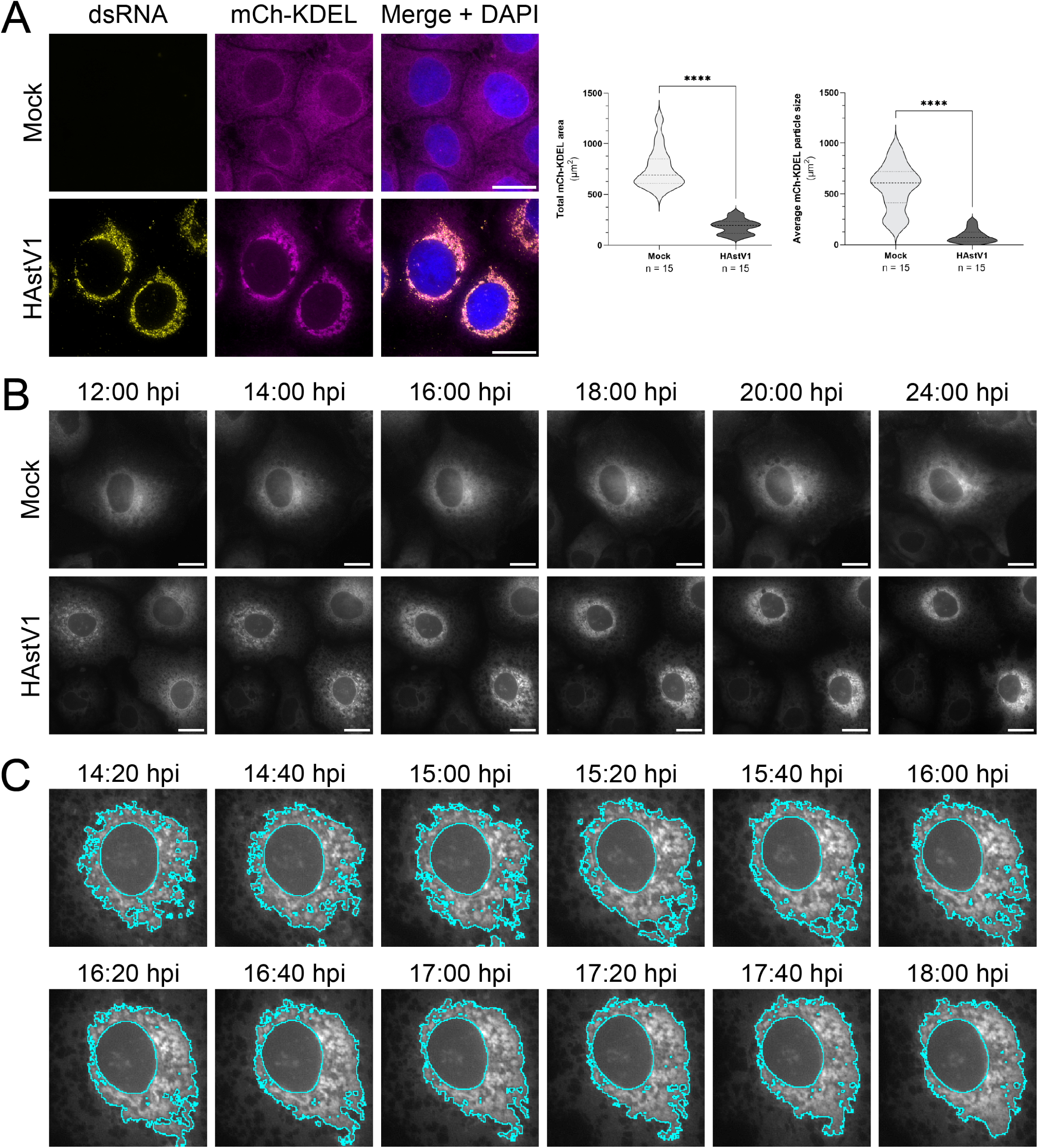
Temporal restructuring of the ER by long term, time-lapse imaging. (A) Representative immunofluorescence staining of mock or HAstV1 infected (MOI = 3; 24 hpi) Huh7 cells expressing the mCherry-KDEL ER marker (magenta). Scale bars represent 20 µm. Quantification of the total area of ER signal (left) and mean area of ER particles (right) between groups was performed by particle analysis using FIJI ImageJ software and an unpaired t-test (**** p < 0.0001). Thresholding was applied uniformly between groups for n = 15 cells across 3 independent experiments. (B) Still frames from live-cell imaging of mock or HAstV1 infected (MOI = 3) Huh7 cells expressing the mCherry-KDEL ER marker (gray) at the indicated times post infection. Scale bars represent 20 µm. (C) A higher magnification of panel B representing images taken between 14:20 and 18:00 hpi. Condensed ER undergoing recruitment is highlighted in cyan.

Ultrastructural transmission electron microscopy (TEM) of HAstV1-infected cells fixed at corresponding time points revealed that DMVs emerge and mature into virion-associated ROs during this same time frame (Fig. 3). Sparse DMVs were observed at 12 hpi and increased in abundance from 14–18 hpi. Further, the distribution of DMVs in large, interconnected networks spanning the perinuclear space of infected cells was most evident at 20 and 24 hpi. Though some individual DMVs were observed, the double membrane of HAstV1-induced DMVs was most frequently comprised of single-membrane vesicles enclosed by an ER-contiguous outer membrane (Fig. 3, black arrows). Additionally, newly formed viral particles showed close association with ER membrane, frequently appearing enclosed within membrane. Together, these findings establish the biogenesis of ER-derived networks of DMVs as a hallmark of HAstV1 infection.

**Figure 3.**
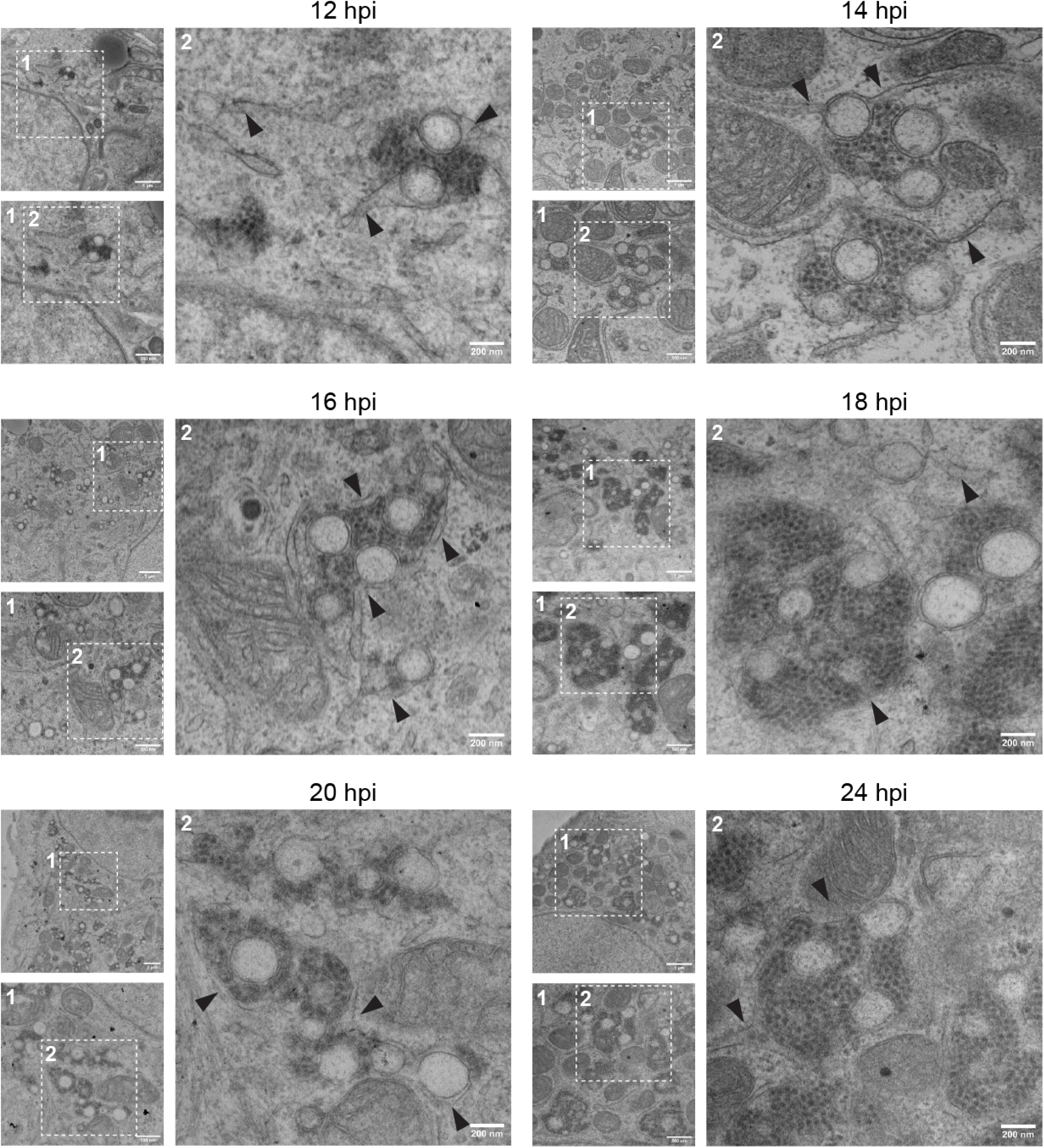
Temporal membrane alterations by transmission electron microscopy. Transmission electron microscopy (TEM) of sectioned HAstV1 infected (MOI = 3) Huh7 cells fixed at the indicated times post infection. Scale bars represent the indicated distances. Insets show a higher magnification of the indicated regions.

### HAstV1 nonstructural proteins colocalize with remodeled ER and replicating RNA

To investigate the contributions of viral nonstructural proteins to RO formation, we generated recombinant viruses expressing epitope-tagged nonstructural proteins to enable localization studies within the context infection-induced ER remodeling. Thus, we inserted a 14-amino acid V5-epitope tag near the N-or C-terminus of each nonstructural protein using our previously published HAstV1 cDNA infectious clone (Fig. S2A) (21). Stable epitope-tagging of VPg and nsp1b was unsuccessful despite rescue of detectable rHAstV1-V5-nsp1b virus. However, recombinant viruses expressing V5-tagged nsp1a/1, nsp1a/2, nsp1a/3, and nsp1a/4 were rescued at similar titers as WT infectious clone-derived virus, suggesting that the epitope tag at the indicated locations does not interfere with genome replication or packaging (Fig. S2A and S2B). To validate the expression of individual V5-tagged nonstructural proteins from infection, we performed immunoblotting of cell lysates infected with each recombinant virus, all of which displayed distinct banding patterns for each expected nonstructural protein (Fig. S2C). We observed bands that corresponded to the predicted sizes of V5-tagged nsp1a/1, nsp1a/2, and nsp1a/3; however, multiple low intensity bands were observed for nsp1a/4. Similar to previous studies that suggested differential processing of nsp1a/4, these findings support the production of functional intermediate proteins (21).

Given that ER-derived DMVs are likely sites of viral replication, we expected HAstV1 viral nonstructural proteins, which are typically involved in replication for +ssRNA viruses, to colocalize with remodeled ER membrane. Thus, we employed our V5-tagged rHAstV1 viruses to investigate the intracellular localization of nonstructural proteins during an active infection. Consistent with this model, nsp1a/1, nsp1a/2, nsp1a/3, and nsp1a/4 colocalized with ER aggregates and replicating RNA (Fig. 4A). Calculation of the Pearson correlation coefficient (PCC) for each group confirmed robust colocalization of all tested nonstructural proteins with mCh-KDEL and dsRNA (Fig. 4B and 4C). Furthermore, the frequent overlap of dsRNA and mCh-KDEL signals supports the role of ER membrane in astrovirus replication (Fig. 4D).

**Figure 4.**
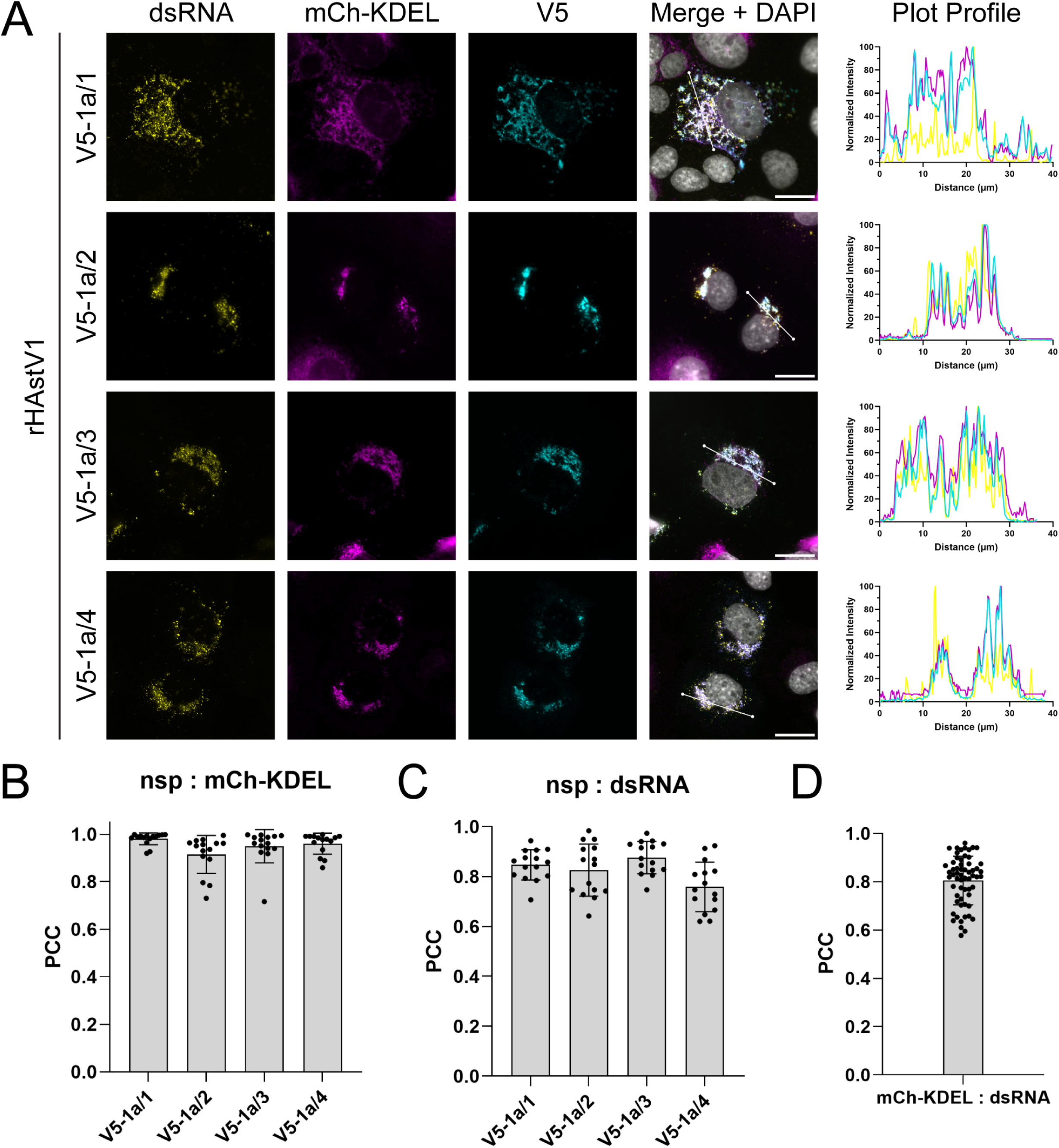
Astrovirus nonstructural proteins colocalize with dsRNA and remodeled ER. (A) Representative immunofluorescence staining of rHAstV1-V5 infected (MOI = 3; 24 hpi) Cos7 cells expressing the mCherry-KDEL ER marker. Percent maximum intensity profiles of dsRNA (yellow), ER (magenta), and V5 (cyan) staining were obtained across a 40 µm linescan shown in merged images using FIJI ImageJ software. (B) Quantification of colocalization of V5-tagged nsps with remodeled ER and (C) dsRNA. Analysis was performed by calculating the Pearson correlation coefficient (PCC) for n = 15 cells from each rHAstV1-V5 group and (D) pooling of all virus-infected groups (total n = 45) to quantify colocalization of dsRNA with remodeled ER across 3 independent experiments.

### HAstV1 replication occurs within the interior of DMVs

The interior compartments of ROs are thought to serve as a secluded niche to protect replicating viral RNA from immune detection (28, 43, 44). However, some +ssRNA viruses assemble their replication complexes on the cytoplasmic face of ROs (45). To elucidate the membrane orientation of HAstV1 replication, we performed IF staining of dsRNA in HAstV1-infected Huh7 cells using a differential permeabilization approach. Selective permeabilization of the plasma membrane with a low concentration of digitonin did not permit antibody-mediated staining of dsRNA, whereas complete permeabilization with Triton X-100 rendered dsRNA epitopes detectable (Fig. 5A and 5B). Similarly, staining of the ER lumen-localized chaperone, calnexin, was only observed after treatment with Triton X-100 (Fig. 5A and 5C). These results indicate that replicating RNA is confined within a membrane-protected compartment, likely the interior of ER-contiguous DMVs.

**Figure 5.**
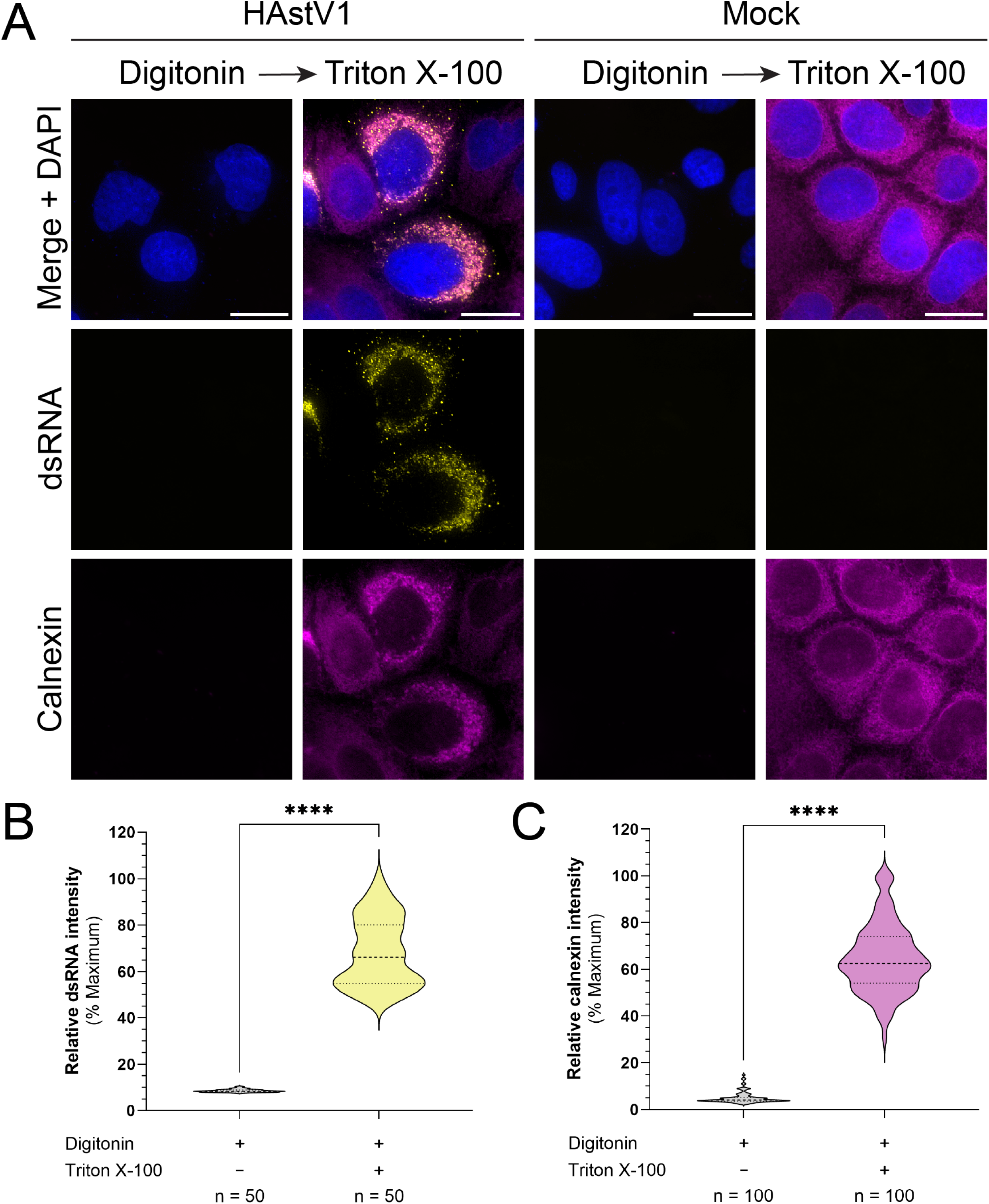
Astrovirus replication occurs within ER-derived DMVs. (A) Representative immunofluorescence staining of mock or HAstV1 infected (MOI = 3; 24 hpi) Huh7 cells permeabilized with digitonin alone or digitonin followed by Triton X-100. Scale bars represent 20 µm. (B) Quantification of the mean fluorescence intensity of dsRNA (yellow) signal for n = 50 cells from HAstV1 infected groups and (C) calnexin (magenta) signal for n = 50 cells from both mock and HAstV1 infected conditions pooled by permeabilization treatment (total n = 100). Mean fluorescence intensity was obtained using FIJI ImageJ software from cells imaged using the same acquisition settings. Data are analyzed by an unpaired t-test and shown as a percentage of the maximum mean intensity observed across three independent experiments (**** p < 0.0001).

### Nsp1a/1 and nsp1a/2 drive ER remodeling and DMV biogenesis

Among other +ssRNA viruses, membrane reorganization is mediated by transmembrane-domain containing nonstructural proteins, where all or the majority of these virus-encoded membrane-interacting proteins coordinate RO biogenesis (28, 46). Notably, astroviruses encode only two nonstructural proteins with transmembrane domains: nsp1a/1 and nsp1a/2. Therefore, we anticipated that these two viral proteins are essential drivers of HAstV-induced ER remodeling. To study the independent roles of these proteins, we generated dual-tagged expression plasmids that encode nsp1a/1, nsp1a/2, or both with an N-terminal green fluorescent protein (GFP) and C-terminal V5-epitope tag. To maintain nsp1a/2 topology, a BiP signal peptide (sp) sequence was added to the N-terminus of GFP (spGFP) to correctly localize the N-terminus of nsp1a/2 to the ER lumen. As expected, expression of either viral protein induced ER remodeling relative to mock and GFP-transfected controls (Fig. 6). Expression of nsp1a/1 formed circular, swollen ER fragments, whereas nsp1a/2 expression resulted in severe fragmentation of the ER spanning the entire cell (Fig. 6A). Co-expression of nsp1a/1 and nsp1a/2 induced ER alterations that most resembled infection; however, quantification of the total area and size of distinct clusters of ER between groups did not resolve these qualitative differences (Fig. 6A).

**Figure 6.**
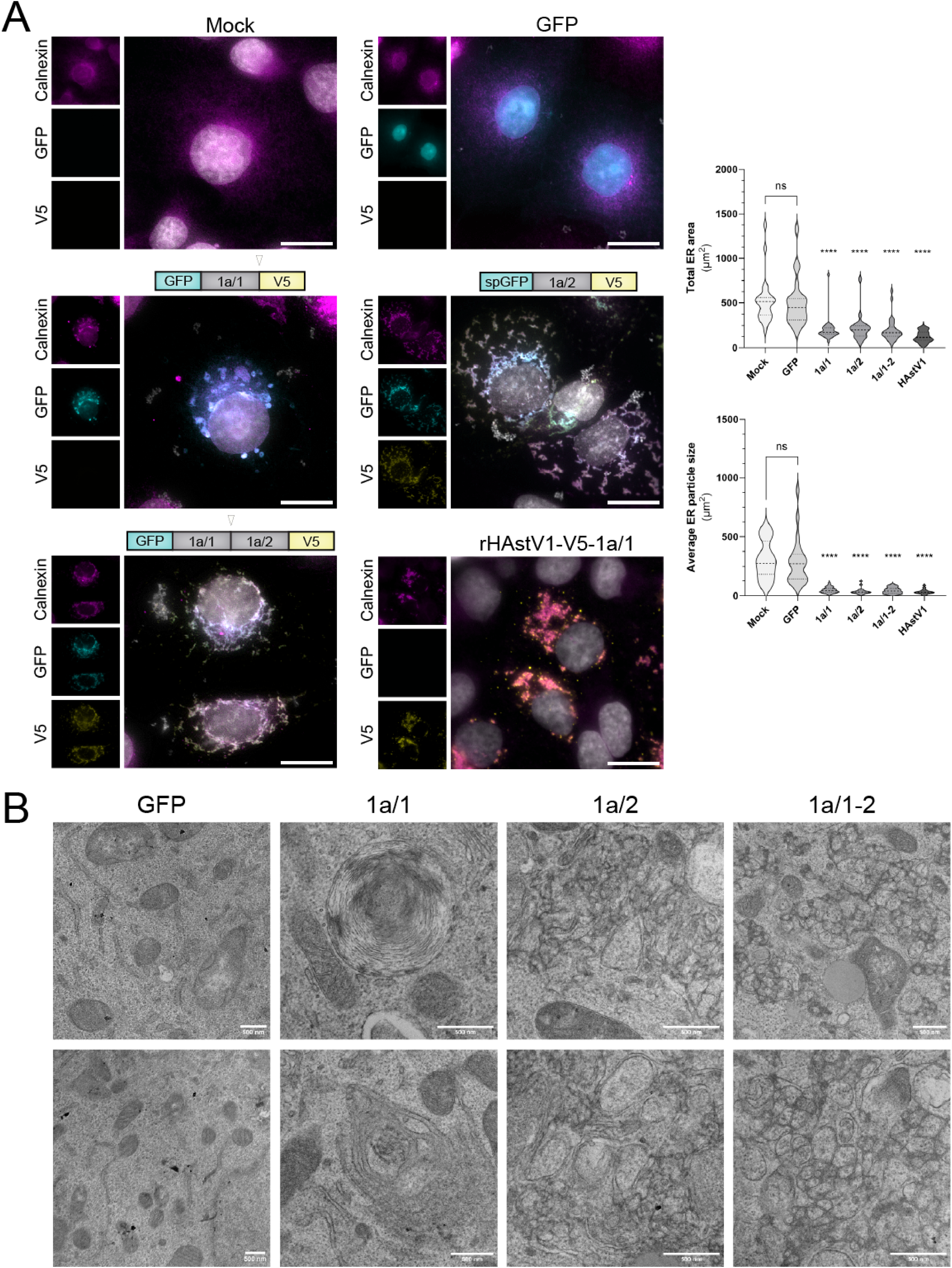
Nsp1a/1 and nsp1a/2 are the minimal constituents for DMV biogenesis. (A) Representative immunofluorescence staining of mock-treated, GFP-transfected, GFP-1a/1-V5 transfected, GFP-1a/2-V5 transfected, GFP-1a/1-1a/2-V5 (1a/1-2) transfected, and rHAstV1-V5-1a/1 infected (MOI = 3) Cos7 cells at 24 hours post treatment. Sites cleaved by host signal peptidase are indicated by white arrowheads. Scale bars represent 20 µm. Quantification of the total area of ER signal (top) and mean area of ER particles (bottom) between groups by particle analysis using FIJI ImageJ software and one-way ANOVA with multiple comparisons versus mock treated cells (**** p < 0.0001). Thresholding was applied uniformly between groups for n = 50 cells across 3 independent experiments. (B) Transmission electron microscopy (TEM) of sectioned Cos7 cells transfected with the indicated constructs and fixed at 24 hours post transfection. Scale bars represent 500 µm.

Further investigation of cellular ultrastructure by TEM confirmed that single expression of each protein resulted in unique membrane alterations, whereas GFP transfection alone did not alter ER morphology (Fig. 6B). We observed the presence of large ER whorls and whorl-like precursors in nsp1a/1-expressing cells. In contrast, disordered networks of convoluted membranes (CM) and high curvature ER were present in nsp1a/2-expressing cells (Fig. 6B). However, only the co-expression of nsp1a/1 and nsp1a/2 (nsp1a/1-2) resulted in the emergence of DMVs resembling infection-induced ROs (Fig. 6B). These nsp1a/1-2-induced DMVs appeared to be tightly packed, likely due to the absence of replicating RNA and assembling viral particles. Furthermore, like infection-induced structures, the DMVs formed from expression of nsp1a/1-2 were linked together by a contiguous outer membrane, forming an expansive network of ER-derived ROs. Together, these findings support a model in which nsp1a/1 and nsp1a/2 are independently capable of ER manipulation, but the coordinated activity between the two is required to establish a functional replication organelle.

### Super resolution dSTORM identifies nsp1a/1 and nsp1a/2 at DMVs

Due to limited spatial resolution, diffraction-limited epifluorescence microscopy is poorly suited to resolve crowded cellular structures. To overcome this limitation, we employed direct stochastic optical reconstruction microscopy (dSTORM) to investigate the nanoscale localization of nsp1a/1 and nsp1a/2 using our V5-tagged nsp1a/1 and nsp1a/2 recombinant viruses.

Infected Huh7 cells displayed large perinuclear ER aggregates captured by particle analysis of clusters exceeding 1 μm^2^, which were absent in all mock-treated cells (Fig. 7A). Furthermore, nsp1a/1 and nsp1a/2 were localized in similar perinuclear aggregates. Notably, the enhanced magnification in this dense region afforded resolution of both nsp1a/1 and nsp1a/2 organized in discrete circular structures with ∼500 nm diameters (Fig. 7B). These structures closely resemble the size and shape of DMVs observed by TEM, which range from 200–500 nm in diameter (Fig. 3) (31). Together, these data support the synergistic roles of transmembrane nonstructural proteins nsp1a/1 and nsp1a/2 as membrane-bound components and functional drivers of DMV biogenesis.

**Figure 7.**
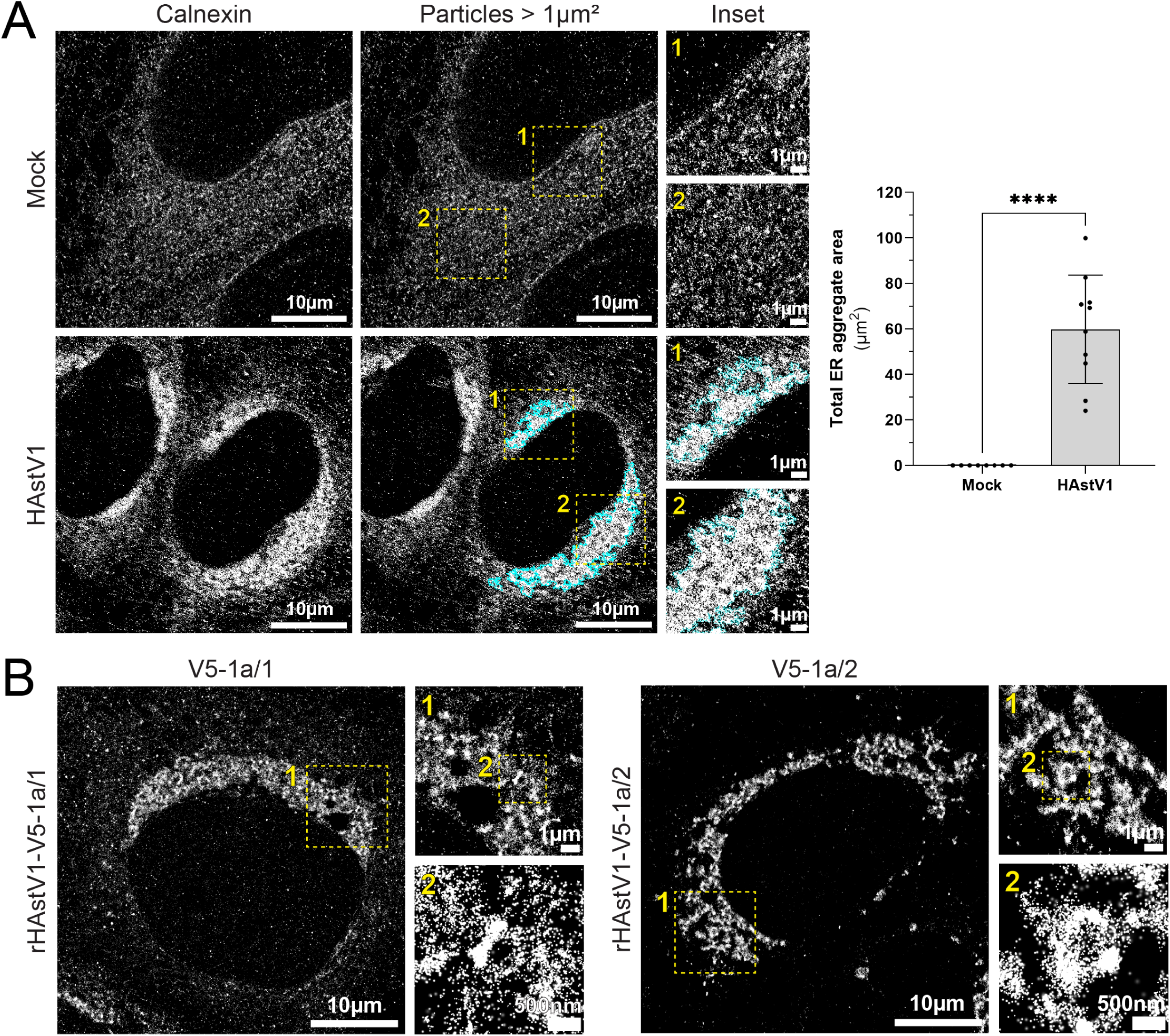
Remodeled ER architecture by super resolution dSTORM. (A) Representative dSTORM images of immunolabeled calnexin in mock or HAstV1 infected (MOI = 3; 24 hpi) Huh7 cells. Segmented particles greater than 1 µm^2^ within a single cell region of interest (ROI) are highlighted in cyan. Insets show a higher magnification of the indicated regions. Scale bars represent the indicated distances. Quantification of the total area of dense ER particles in mock and HAstV1 infected cells was performed by thresholding and particle analysis using FIJI ImageJ software and an unpaired t-test (*** p < 0.001). (B) Representative dSTORM images of immunolabeled V5 in rHAstV1-V5-1a/1 and 1a/2 infected (MOI = 3; 24 hpi) Huh7 cells. Insets show a higher magnification of the indicated regions. Scale bars represent the indicated distances.

## Discussion

In this study, we demonstrate vast remodeling of the ER driven by viral nsp1a/1 and nsp1a/2 during HAstV1 infection (Fig. 8). Our results indicate that these proteins act synergistically to drive the formation of large ER-derived DMV networks that support virus RNA replication and virion production. The ER is a highly dynamic organelle composed of a contiguous membrane that plays a central role in cellular homeostasis and biosynthetic processes, such as protein translation and folding, lipid biosynthesis and glycosylation, and secretion (42, 47–49). Due to its significance in mediating host processes, the ER is a frequent target of +ssRNA viruses, many of which remodel ER membrane to generate specialized compartments for replication (43, 44). These ROs are essential hubs of viral RNA synthesis that optimize host and viral protein-protein and protein-RNA interactions and segregate replication intermediates from immune sensors present in the cytosol (46, 50, 51).

**Figure 8.**
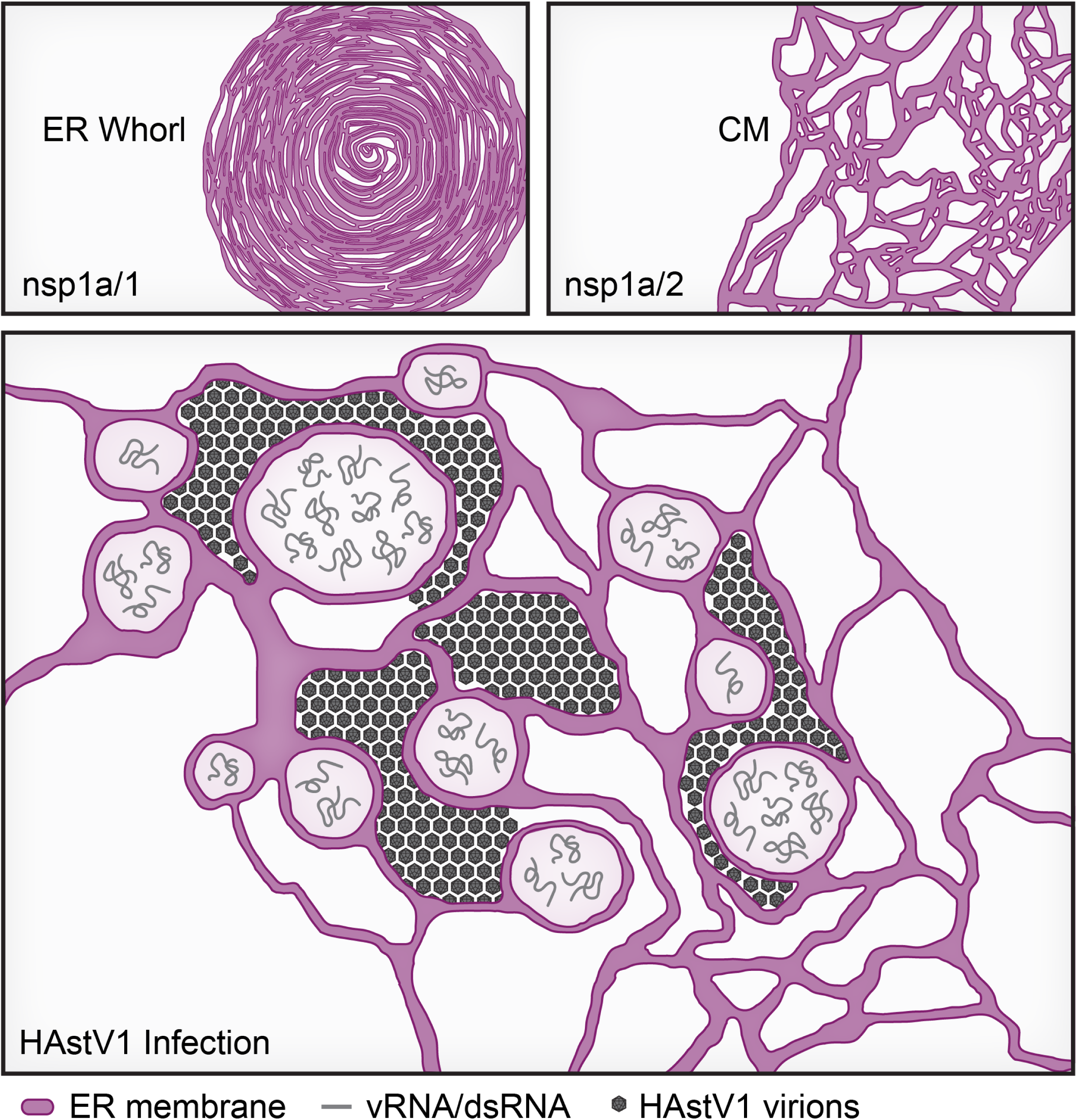
Proposed model of HAstV-induced membrane reorganization. Expression of the single-pass transmembrane viral protein nsp1a/1 led to the formation of ER whorls, whereas the expression of the multi-pass transmembrane viral protein nsp1a/2 resulted in the emergence of convoluted ER membranes (CM). These two ER-anchored nonstructural proteins function synergistically during infection to drive the biogenesis of HAstV ROs, which contain replicating RNA and coordinate virus assembly sites.

Among different +ssRNA viral families, remodeling of host membranes is driven by nonstructural proteins with transmembrane or membrane-tethering domains. For example, dengue virus (DENV) generates replicase-containing vesicle packets that begin as dilatated ER sheets and progressively invaginate to enclose vesicles (52–55). Expression of the highly hydrophobic NS4A and NS4B proteins of West Nile virus and DENV has been shown to drastically alter ER architecture (34, 56–58). In contrast to orthoflavivirus ER invaginations, hepatitis C virus (HCV) infection induces the formation of DMVs by ER ex-vaginations, where the outer membrane of the DMV remains contiguous with the ER by a short stalk that can detach to yield singular DMVs (59). During late stages of HCV infection, a variety of membranous structures emerge, including double membrane tubules (DMTs) and multi-membrane vesicles, formed by interactions involving NS3–NS5A (59–61). Coronaviruses (CoV), including SARS and MERS CoVs, induce the accumulation of DMVs formed by nsp3, nsp4, and nsp6 (39, 40, 46, 62). Like HCV DMVs, CoV DMVs are often linked by an ER-contiguous outer membrane and a network of convoluted membranes (CM) (63). The co-expression of nsp3 and nsp4 is sufficient to generate DMVs, whereas nsp6 is responsible for establishing ER connectors by oligomerization and membrane zippering activity (64, 65). Visually, astrovirus DMVs most closely resemble CoV ROs, with an ER-contiguous outer membrane often linking multiple vesicle clusters. However, the membrane source and viral proteins involved in astrovirus RO biogenesis have yet to be experimentally characterized (31). Based on colocalization of several viral proteins with the ER and perinuclear anchoring of replication complexes, astrovirus ROs have previously been assumed to be ER-derived (23, 26, 31).

To investigate the nonstructural proteins contributing to HAstV1 membrane rearrangement, we utilized our HAstV1 reverse genetics system to create recombinant viruses expressing V5-epitope-tagged nonstructural proteins (21). Consistent with previous studies, we demonstrated that infection resulted in colocalization of viral nonstructural proteins with dsRNA and remodeled ER membrane (23, 26). Immunoblots of infected cell lysates probing for the V5-epitope tag confirmed the production of V5-tagged nonstructural proteins that corresponded to previously predicted protein sizes (21). However, these results also revealed several previously uncharacterized intermediate bands, as well as differences in cleavage efficiency among final products (Fig. S2C). With the recent identification of the viral protease cleavage motif, further studies are now possible to identify the importance of these intermediate bands (21).

Virus-induced membrane alterations serve as isolated environments that shield viral RNA from host immune surveillance (28, 43, 44). Nevertheless, some picornaviruses replicate in association with the cytoplasmic surface of ROs (45). To determine the site of RNA synthesis for HAstV1 ROs, we performed IF staining of dsRNA using a low concentration of digitonin to selectively permeabilize the plasma membrane. Using this strategy, we determined that dsRNA is not exposed to the cytosol, suggesting that HAstV1 replication is confined to the interior of infection-induced DMVs. For viruses that replicate within enclosed ROs, the translation of structural proteins occurs in a topologically separate environment from replication. Thus, these viruses devise specialized strategies to allow RNA to reach the cytoplasm for packaging into progeny viral particles. Though the exact membrane orientation of HCV replication is unclear, both HCV and orthoflavivirus ROs have been observed to contain small openings to the cytoplasm that could accommodate release of positive sense RNA for packaging (54, 59).

However, the existence of viral or host proteins regulating this process has not been characterized. In contrast, recent studies have identified a CoV pore complex consisting of nsp3 and nsp4 that bridges the DMV interior to the cytoplasm, enabling viral RNA export and capture by nucleocapsid proteins (66–68). Given the membrane orientation of HAstV1 replication, it is possible that astroviruses may employ a similar strategy to manage virion assembly. Future studies leveraging structural biology and reverse genetics approaches may uncover the mechanisms of astrovirus RNA packaging.

As infection progresses, ROs are thought to mature to address the transitional needs of the virus, such as spatial-temporal regulation of replication and packaging (69–73). In support of this, assembled viral particles were found directly adjacent to DMVs, with an abundance of packaged virus found at later stages of infection (Fig. 3). Consistent with studies demonstrating a membrane affinity for astrovirus virions, we frequently observed viral particles associated with or encircled by ER membrane (74, 75). These findings may suggest a role for capsid in the formation of ER-associated virus assembly sites. Membranous “bags” of viral particles have been reported among enveloped viruses that assemble at the ER and ER-golgi intermediate complex (ERGIC); however, this is an uncommon feature among non-enveloped viruses like astroviruses (54, 76, 77). Some enteroviruses, such as poliovirus and enterovirus 71, can utilize an alternative egress strategy involving autophagosome-mediated exit without lysis (AWOL) that results in release of viral particles in exosomes (78–81). Though extracellular vesicles containing astrovirus particles have not been reported, astrovirus egress is characteristically non-lytic, suggesting that membrane association of virions could aid in virus release (75, 82).

Notably, some host components involved in the initiation of autophagy, such as the phosphatidylinositol 3-kinase (PI3K) complex, have been shown to support astrovirus replication and DMV formation, underscoring the importance of autophagy in astrovirus morphogenesis (31). However, the current literature highlights a lack of knowledge surrounding astrovirus egress; thus, the mechanisms by which HAstVs coordinate virion assembly, trafficking, and release from infected cells remain unclear.

In this study, we identified that nsp1a/1 and nsp1a/2 are functional drivers of RO biogenesis and structural components of DMV membranes. The expression of nsp1a/1 and nsp1a/2 individually induced extensive restructuring of the ER, but the presence of both viral proteins was required to form DMVs that closely resemble infection-induced membrane alterations. However, the underlying contributions of these proteins RO structure and function remain elusive. Nsp1a/1 contains a single transmembrane domain that also functions as a signal peptide to direct translation of viral polyproteins to the ER (1, 23). As such, it is removed from the N-terminus of polyproteins through a highly efficient cleavage event performed by a host signal peptidase (22, 23). Previous studies demonstrated that conserved di-arginine motifs present within nsp1a/1 contribute to the perinuclear retention of replication complexes, as alanine substitution of these motifs resulted in diffuse ER staining of nsp1a/1 (23). Consistent with these findings, we observed nsp1a/1 localized to clustered ER membrane. However, further investigation by super resolution dSTORM revealed the presence of a small proportion of nsp1a/1, but not nsp1a/2, dispersed in a reticular network that was not resolvable by traditional epifluorescence imaging (Fig. 7B). Similar to observations made by IF and live-cell imaging, there is often weak-staining of ER membrane in the periphery of infected cells at 24 hpi (Fig. 1 and 2). Together, these data suggest that diffusely localized nsp1a/1 may contribute to the recruitment of ER membrane to perinuclear aggregates, perhaps by coordination of di-arginine motifs, or by an otherwise uncharacterized mechanism. Further, ectopic expression of nsp1a/1 resulted in the accumulation of extensive ER whorls observed by TEM, structures formed in response to ER stress (83). Thus, it remains unclear whether the contribution of nsp1a/1 to DMV biogenesis is by direct manipulation of ER membrane or by the induction of ER stress.

Though its specific membrane topology is poorly understood, the nsp1a/2 protein of human astroviruses is predicted to contain 4–6 transmembrane domains (1). Furthermore, the functional significance of nsp1a/2 during infection remains elusive. In this report, we demonstrated that expression of nsp1a/2 induces substantial fragmentation of the ER network and the formation of extensively convoluted ER membrane (Fig 6). In contrast to nsp1a/1, dSTORM of cells infected with a V5-nsp1a/2 encoding virus revealed that nsp1a/2 localizes exclusively to perinuclear aggregates corresponding to remodeled ER, suggesting that its role is specific to DMVs and the membranes altered for DMV biogenesis (Fig 7B). Among +ssRNA viruses such as orthoflaviviruses, coronaviruses, and picornaviruses, highly hydrophobic, multi-pass transmembrane proteins are commonly implicated in virus-induced membrane alterations (28). Thus, we establish nsp1a/2 as a previously unrecognized determinant of ER-derived RO biogenesis during HAstV1 infection.

Collectively, this study revealed that restructuring of ER membrane into DMV networks is a defining feature of productive HAstV1 infection. Furthermore, we identified the two viral proteins responsible for ER manipulation and DMV formation: nsp1a/1 and nsp1a/2. Future studies will be directed at elucidating the specific mechanisms of astrovirus RO formation by these two viral proteins. Thus, our findings will provide insights into the rational design of nsp1a/1 or nsp1a/2-targeted inhibitors, which could serve as effective antivirals to limit astrovirus replication.

## Materials and Methods

### Cell culture

Caco2 cells (ATCC, HTB-37) were maintained in modified Eagle’s medium supplemented with 20% FBS, 10% sodium pyruvate, 10% non-essential amino acids and 100 IU penicillin per 100 μg/mL streptomycin. Human hepatoma cells (Huh7) were maintained in Dulbecco’s modified Eagle’s medium supplemented with 10% FBS and 100 IU penicillin per 100 μg/mL streptomycin and 9 g/L glucose. Cos7 cells (a gift from Dr. Alexa Mattheyses, University of Alabama at Birmingham) and HEK293T cells (ATCC, CRL-11268) were maintained in Dulbecco’s modified Eagle’s medium supplemented with 10% FBS and 100 IU penicillin per 100 μg/mL streptomycin.

Hybridoma cell lines (ATCC, CRL-8795) that were used to generate monoclonal antibody 8E7 (mouse anti-HAstV1 capsid protein) were grown in Iscove’s modified Dulbecco’s medium with 2mM L-glutamine and 15% heat inactivated fetal bovine serum. All cells were cultured at 37°C and 5% CO_2_.

### Preparation of virus stocks

Virus was recovered by transfection of Huh7 cells with pcDNA-HAstV1 or pcDNA-HAstV1-V5 plasmids using polyethylenimine (PEI, 25 kDa) at a 1:1 ratio of DNA (μg) to 1 mg/mL PEI stock. At 12 hours post transfection, cells were gently washed with PBS, and the media was replaced with serum-free media containing 10 μg/mL porcine trypsin. At 72 hours post transfection, cells were lysed in their supernatants by three subsequent freezes in liquid nitrogen and thaws at 37°C. Cell debris was removed by centrifugation at 5,000 x g for 5 minutes, then supernatants were collected and stored at −80°C until use.

### Virus titration

Caco2 cells were seeded in a 96-well plate at 2.5 × 10^4^ cells/well and allowed to grow for 72 hours to reach confluence. Media was replaced with serum-free Caco2 media supplemented with 0.3% bovine serum albumin (BSA) for an hour, followed by infection with 10-fold dilutions of virus-containing supernatant. At 20 hours post infection, cells were fixed in PBS + 4% paraformaldehyde (PFA), permeabilized with PBS + 0.1% Triton X-100, then washed with PBS. Primary incubation with mouse anti-HAstV1 capsid protein (8E7) diluted 1:4 in PBS was performed overnight at 4°C. Following primary, cells were washed with PBS, then incubated goat anti-mouse Alexa Fluor 488-conjugated secondary antibody diluted 1:1000 in PBS for 30 minutes, followed by two final PBS washes. Capsid positive foci were counted in technical duplicate using an Olympus IX83 inverted fluorescence microscope to calculate titer (FFU/mL).

### Immunofluorescence microscopy

Cells were grown in 8-well chamber slides (Celltreat), fixed in PBS + 4% paraformaldehyde (PFA), permeabilized with PBS + 0.1% Triton X-100, then washed with PBS. Primary incubation was performed overnight at 4°C with antibodies diluted in PBS. Primary antibodies used for immunofluorescence were as follows: rabbit anti-calnexin (Abcam, ab22595, 1:1000), mouse anti-dsRNA (Kerafast, 1:25), mouse anti-V5 (Invitrogen, 46-0705, 1:5000), rabbit anti-V5 (Cell Signaling, 13202, 1:1000), and mouse anti-HAstV1 capsid (8E7, 1:4). Following primary incubation, cells were washed with PBS, then incubated with corresponding Alexa Fluor 488, 594, or 647-conjugated secondary antibody diluted 1:1000 in PBS for 30 minutes, followed by a PBS wash. Cells were incubated in 300nM 4’,6-diamidino-2-phenylindole (DAPI) for 5 minutes, followed by two final PBS washes. Glass slides were mounted using a glass coverslip and ProLong Diamond Antifade Mountant (Invitrogen, P36965), then allowed to cure overnight at room temperature. Epifluorescence images were acquired using an Olympus IX83 inverted fluorescence microscope.

### Immunoblots

Infected Caco2 cells were lysed in a 1X RIPA + protease inhibitor cocktail (Sigma) at 24 hours post infection. Lysates were clarified by centrifugation at 12,000 x g and 4°C for 10 minutes, then prepared with a 6X SDS loading buffer + betamercaptoethanol (BME). Prepared lysates were separated by SDS-PAGE on a 4-20% Tris-glycine polyacrylamide pre-cast gel (BioRad) for 50 minutes, followed by transfer to nitrocellulose membrane for 50 minutes. Membranes were blocked for 30 minutes in PBS + 10% non-fat milk, then probed using primary antibodies diluted in PBST + 5% BSA. Primary antibodies used include mouse anti-V5 (Invitrogen, 1:5000), rabbit anti-GAPDH (ProteinTech, 1:2000), rabbit anti-Calnexin (GeneTex, 1:5000), rabbit anti-CKAP4/CLIMP63 (ProteinTech, 1:5000), rabbit anti-RTN4 (ProteinTech: 1:5000). Proteins were observed using near-infrared dye-conjugated secondary antibodies (LiCor) diluted in PBST + 5% non-fat milk, then imaged on a LiCor Odyssey CLx imaging system.

### Plasmids

All pcDNA-HAstV1-V5 tagged infectious clone plasmids were cloned by assembly of 5’ and 3’ PCR fragments into the PstI and AgeI restriction sites of pcDNA-HAstV1 (21). For all reactions, the vector was generated by digesting pcDNA-HAstV1 with restriction enzymes PstI and AgeI (New England BioLabs). The 5’ PCR fragment was generated using Q5 High Fidelity DNA polymerase (New England BioLabs) with AstV-PstI_F and a site-specific reverse primer. Site specific reverse primers used were V5-1a/1_R, V5-1a/2_R, V5-1a/3_R, and V5-1a/4_R (Table S1). The 3’ PCR fragment was generated with a site-specific forward primer and AstV-AgeI_R. Site-specific forward primers used were V5-1a/1_F, V5-1a/2_F, V5-1a/3_F, and V5-1a/4_F (Table S1).

The generation of pcDNA-GFP-V5, used as a vector for assembly of dual-tagged viral protein expression plasmids, was performed as previously described (21). The pcDNA-GFP-V5 plasmid was linearized with BamHI (New England BioLabs) and assembled with DNA encoding viral proteins. PCR products for the generation of GFP-1a/1-V5, GFP-1a/2-V5, and GFP-1a/1-2-V5 were amplified from pcDNA-HAstV1 using Q5 High Fidelity DNA polymerase and the following respective primers: 1a/1_F with 1a/1_R, 1a/2_F with 1a/2_R, and 1a/1_F with 1a/2_R (Table S1). Generation of spGFP-1a/2-V5 was accomplished by assembly of pcDNA-GFP-V5 digested with BamHI and MluI (New England BioLabs) with two PCR fragments amplified from GFP-1a/2-V5 using the following respective primers: MluI-CMV_F with SpBIP-GFP_R, and SpBIP-GFP_F with 1a/2_R (Table S1).

All PCR products and vectors were purified by agarose gel electrophoresis and extracted with Quick Gel extraction kit (Invitrogen, K220010). Plasmids were assembled by HiFi Assembly (New England BioLabs), and reactions were transformed into Stable Competent *E. coli* (New England BioLabs, C3040I) at 30°C. Plasmids were purified by ZymoPURE Express Plasmid Midiprep Kit according to the manufacturer’s protocol, then sequenced using Plasmidsaurus.

### Transfections and lentivirus production

Cos7 cells were transfected using TransIT LT-1 transfection reagent (Mirus Bio) according to the manufacturer’s protocol. For generation of mCherry-KDEL lentivirus, HEK293T cells were transfected with 1 µg pLJM1-mCherry-KDEL, 0.75 µg psPAX (a gift from Didier Trono, Addgene plasmid #12260), and 0.25 µg pCAGGS-G-Kan (a gift from Todd Green, University of Alabama at Birmingham) using polyethylenimine (PEI, 25 kDa) at a 1:1 ratio of DNA (μg) to 1 mg/mL PEI stock. Cells were incubated at 37°C for 48 hours, then cell supernatant was harvested. Debris was pelleted by centrifugation at 300 x g for 5 minutes, then lentivirus-containing supernatants were collected and stored at −80°C until further use.

### Live-cell imaging

Huh7 cells were transduced with pLJM1-mCherry-KDEL lentivirus at 1 × 10^4^ cells/well in an 8-well polymer live imaging slide (Ibidi), then allowed to grow for 48 hours. Media was removed and replaced with virus at an MOI of 3 diluted in serum free media. After absorption for 1 hour at 37°C, well volume was brought up to 200 μL with serum-containing media. Live cell imaging was performed using an Olympus IX83 inverted fluorescent microscope in a humidified chamber at 37°C and 5% CO_2_. Epifluorescence images were acquired every 20 minutes for 20 hours using the autofocus function. Final images at 24 hpi were acquired manually using the same exposure settings.

### Transmission electron microscopy

Cos7 and Huh7 cells were seeded at 4 × 10^5^ cells/well in a 6-well plate and allowed to adhere for 24 hours prior to transfection or infection. Cells were fixed using 3% glutaraldehyde in 0.1M cacodylate buffer for 1 hour, then scraped and pelleted at 300 x g for 5 minutes to preserve ultrastructure. Carbon TEM grids overlaid with thin slices of sectioned cell pellets were prepared at the University of Alabama at Birmingham High Resolution Imaging Facility. Imaging was performed using a JOEL 1400 HC Flash TEM at 120 kV with an AMT NanoSprint43 Mk-II CMOS camera.

### Super resolution dSTORM

Samples were prepared as described for immunofluorescence microscopy in an 8-well glass bottom live-imaging slide (Ibidi) using Alexa Fluor 647-conjuagated secondary antibodies.

Imaging was performed in a photoswitching buffer containing 50mM Tris-HCl and 10mM NaCl supplemented with 10% (w/v) glucose, 2% catalase and glucose oxidase solution (GLOX) and 50mM cysteamine (MEA) (a gift from Dr. Alexa Mattheyses, University of Alabama at Birmingham). dSTORM images were obtained using a Nikon Ti-2 microscope using a 100X 1.49 NA oil immersion objective and 647 nm excitation laser at the University of Alabama at Birmingham High Resolution Imaging Facility. For each image, 10,000 frames were acquired and reconstructed in Nikon Elements.

### Image and statistical analysis

Image analysis was performed using FIJI ImageJ software with statistical analysis in GraphPad Prism 10. For analysis of epifluorescence ER morphology, signal overlapping with DAPI was excluded to eliminate ER signal corresponding to out of focus light from thick cells. The signal intensity histogram was scaled to fit values of 0 to 255, and thresholding was performed under the same conditions between mock and HAstV1 infected cells to capture signal for particle analysis in single-cell regions of interest (ROI) drawn using the freehand selection tool. Particles less than 2 µm^2^ were excluded due the presence of pixelated edges from thresholding. For analysis of mean fluorescence intensity between digitonin and fully permeabilized cells, all images across three independent replicates were acquired using the same exposure settings.

Mean fluorescence intensity from raw images was captured in single-cell ROI, then transformed as a percentage of the maximum signal. For analysis of dSTORM ER morphology, reconstructed images were exported at 4 nm per pixel and subjected to Gaussian blur preprocessing (σ = 2) to merge closely spaced pixels within dense localization clusters.

Thresholding was performed with no exclusion criteria to capture total signal for particle size analysis in single-cell ROI. The total area of particles greater than 1 µm^2^ was measured to quantify regions of dense ER aggregation and exclude diffuse ER staining. Figures were generated using Adobe Illustrator.

## Acknowledgements.

This research was supported by National Institutes of Health R35-GM150638 (N.J.L.) and T32-GM146611 (B.B.), as well as the University of Alabama at Birmingham Heersink School of Medicine (N.J.L.). Research reported in this publication was supported by the UAB High Resolution Imaging Facility.

